# Transcriptional feedback of Erk signaling waves in zebrafish scale regeneration

**DOI:** 10.1101/2025.11.19.689292

**Authors:** Coline Coudeville, Tristan Guyomar, Julie Allamand, Franka Voigt, Alessandro De Simone

## Abstract

Regeneration requires long-range cellular coordination. In regenerating bony zebrafish scales, waves of activity of the Extracellular Receptor Related Kinase (Erk) induce growth of bone-forming tissue. Erk waves were proposed to result from an excitable system including feedback with Erk activators and inhibitors. Here, we characterize how Erk modulates its inhibitors, predicted to limit wave frequency and thus tissue growth. First, we found that Erk waves can form spirals, signature of excitable systems, and exhibit refractory behavior compatible with Erk inhibitors following Erk waves. Using a newly developed single-molecule fluorescence in situ hybridization (smFISH) technique for whole scales, we discovered that Erk waves induce trailing transcript waves of the inhibitors *dusp5, spry2* and *spry4* through Rectified Linear Unit (ReLU) responses. Furthermore, we discovered that Erk waves modulate transcription of the gene *osterix*, which controls maturation of bone-forming cells. This reveals a transcriptionally-encoded mechanism that couples Erk wave dynamics, and thus tissue growth, with scale maturation during regeneration.

## Introduction

During regeneration, cells coordinate with each other to rebuild a body part of the right size and shape. In vertebrate appendages, tissue regeneration involves thousands of cells that communicate through diverse signaling inputs (*1-3*). These “morphogenetic inputs” have been classically described as extracellular diffusible ligands activating signaling pathways, but mechanical and electric signals have also been implied in size and shape control in various systems (*3*). One critical coordination mechanism relies on the signaling pathway Extracellular Receptor Related Kinase (Erk). Quantitative studies of Erk activity allowed to discover that the Erk pathway activates transiently in long-range travelling waves across homeostatic, healing and regenerating tissues (*4-9*). The properties of these Erk activity waves (in short, Erk waves) and their effects vary between systems. For example, in amputated planaria, a peak of Erk activity travels across the entire flatworm at a speed of hundreds of cells per hours, delivering information that initiates regeneration (*9*). In wounded epithelia, including in mice, a wave of Erk activation and cell contractility propagates away from the wound site at a speed of roughly ten cells per hour and triggers cell migration toward the wound itself (*4, 5, 10, 11*). In bony zebrafish regenerating scales, Erk activates in circular and expanding ring-like patterns that move at a much lower speed, in the order of one cell per hour (*6*). In these scale waves, Erk stays active in the same cell for many hours and triggers localized cell growth without cell division (hypertrophy). The more waves travel across the tissue, the more the scale grows. Therefore, the frequency of wave generation controls the rate of scale growth. While details regarding each of these waves differ between systems, in all cases they behave as transient peaks of Erk activity moving across space. Despite these discoveries, it remains unclear what is the mechanism that controls the properties of Erk waves, as well as how they control tissue dynamics. In scales, wave frequency was proposed to depend on Erk inhibitors, activated by Erk itself, which would switch Erk off after activation, and delay further wave formation (*6, 12*). In this work, we report that, in scales, Erk waves induce transcriptional waves of their own inhibition machinery, as well as of bone cells differentiation, thus coupling wave dynamics and bone maturation with scale growth.

Zebrafish scales are disk-shaped skin appendages, formed by an acellular millimeter-sized bone disk covered by a monolayer of bone-forming cells (Fig. 1A, B) (*13-17*). Cells of this bone-forming monolayer share functional and molecular characteristic with mammalian bone-depositing osteoblasts (*14, 15, 17, 18*) and, therefore, have been referred to as “osteoblasts”. After scale loss and regeneration onset, a new pool of osteoblasts forms, likely by dermal differentiation, followed by cell proliferation (*15*). Eventually, around the 4^th^ day of regeneration, proliferation stops and tissue growth continues by osteoblast hypertrophy (*15*). Live imaging of an Erk biosensor, quantifications and theory, together, revealed that osteoblast hypertrophy is induced by travelling waves of Erk activity that start centrally and propagate outward as expanding rings (Fig. 1C, D) (*6*). These results were obtained using a transgenic Erk biosensor (Erk Kinase Translocation Reporter – Erk KTR; Fig. S1A) (*6, 19*). The fluorescently tagged Erk KTR contains an Erk binding site and nearby phosphorylation sites; phosphorylation causes the Erk KTR to translocate from the nucleus to the cytoplasm. Therefore, the ratio of Erk KTR fluorescence in the cytoplasm and in the nucleus is a proxy of Erk activity.

**Figure 1.**
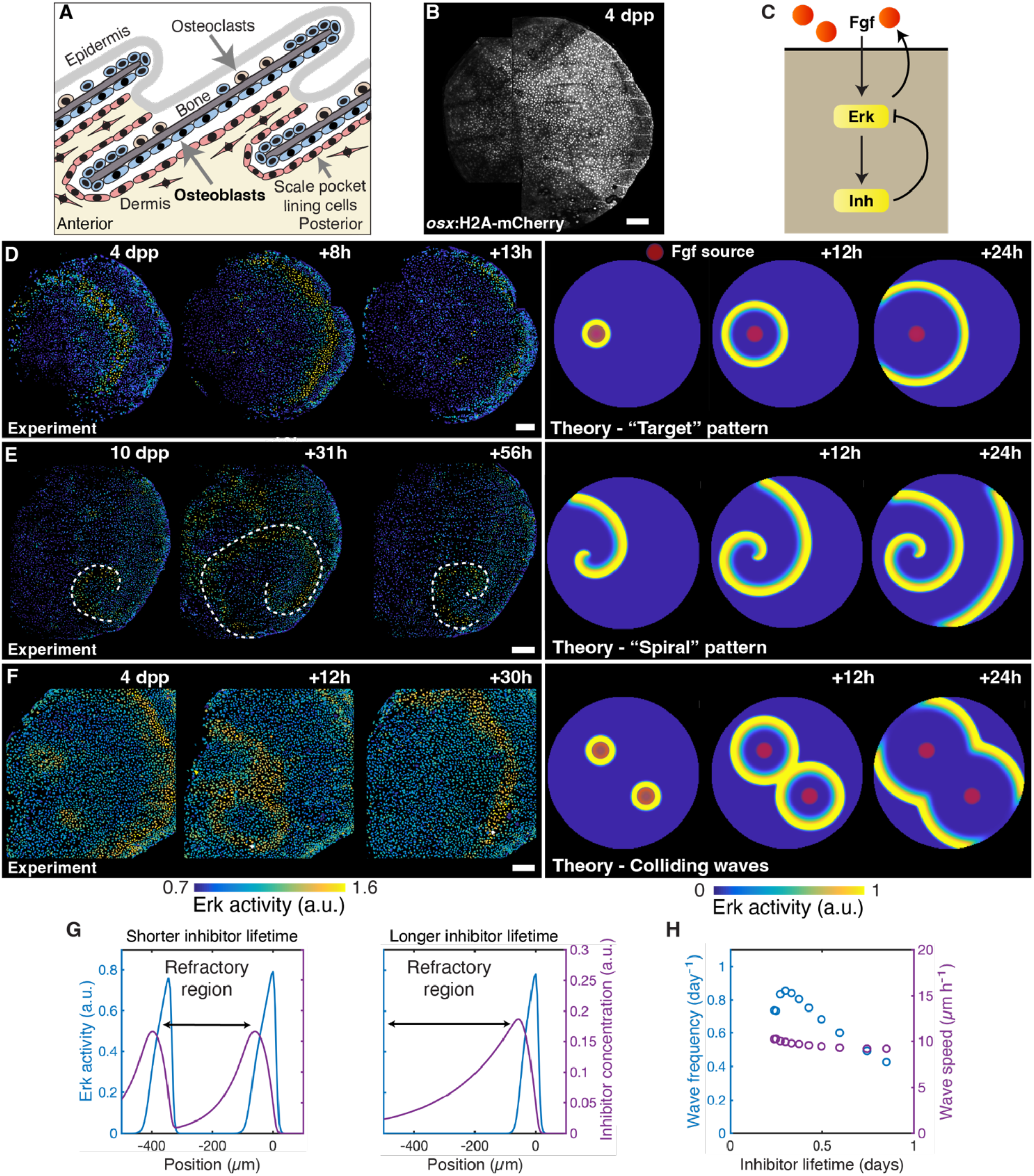
Excitability of Erk activity waves in zebrafish scale regeneration. A. Schematics of scale anatomy (depicted in cross-section). B. Example of osteoblast nuclei in a regenerating scale (nuclear marker: osx:H2A-mCherry). C. Model of excitable Erk activity waves, including a positive feedback loop between Erk and a diffusible Erk activator (Fgf) and a negative feedback loop between Erk and a non-diffusible Erk inhibitor (arrows: activating interactions; blunt arrows: inhibiting interactions). D. Example of an Erk activity wave in vivo (left; n> 30) and in theoretical simulations (right - red shaded region: constant source of activator). E. Example of a spiral Erk activity wave in vivo (left; dashed line: spiral wavefront; a second spiral wave was detected in the same scale at later time-points; n = 2, see Fig. S1B) and in theoretical simulations (right). F. Example of colliding waves in vivo (left; n = 9 from this work and a re-analyzed dataset from (6); a subset of scales was treated with DMSO) and theoretical simulations (right - red shaded region: constant source of activator). G. Erk activity and inhibitor concentration profiles in theoretical simulations with two different lifetimes of the Erk inhibitor. Erk activity waves are predicted to be followed by a region of high inhibitor concentration. H. Theoretical prediction of wave frequency and speed as a function of the lifetime of the Erk inhibitor. a.u.: arbitrary unit. dpp: days post-plucking. Scale bars, 100 µm. Panel A and C is adapted from (6); Copyright © 2021, The Author(s), under exclusive license to Springer Nature Limited.

Experimental work using the Erk KTR biosensor, together with theory, indicated that Erk waves could be generated by an excitable system (*6*). An excitable system exhibits a large oscillation from its resting state following a sufficiently large stimulus, before returning to this resting state (*20, 21*). This behavior was proposed to result from a positive feedback loop between Erk and Fibroblast Growth Factors (Fgfs), and a negative feedback loop between Erk and its inhibitors (*6*). Theory that includes negative feedback loops between Erk, Fgfs and Erk inhibitors, together with a localized source of Fgf, reproduces a phenomenology that is similar to the one observed in experiments (Fig. 1C, D, Movie S1) (*6*). Theory predicts that the positive feedback loops controls wave speed, while the negative feedback loop limits wave frequency (*12*). Typically, in simulations, Erk forms a “target pattern” in which a ring-like Erk wave propagates away its source (Fig. 1D, Movie S1). However, in some conditions, excitable systems can also form characteristic self-sustained spirals (*20-22*). Spiral patterns have been observed experimentally in various systems, including chemical reactions and cardiac electrophysiology (*21*). Spirals can also be formed in the model of Erk activity propagation (Fig. 1E Theory, Movie S2), but we typically observed circular target patterns in experiments (N> 100 scales). Serendipitously, we recently observed a spiral starting from the posterior of a regenerating scale, repeated after some time in the same location (Fig. 1E Experiment, Movie S3). Prompted by this observation, we re-analyzed current and past datasets and observed a second instance of a spiral pattern (Fig. S1B). While non-characteristic of Erk signaling in scale regeneration, the fact that this behavior is possible is a signature of excitable dynamics.

The theory of Erk activity waves excitability posits that Erk stimulates its own inhibitors, which are responsible for switching Erk off (*6*). Indeed, bulk RNA sequencing of a cellular pool enriched in Erk active osteoblasts showed higher levels of transcripts of the Erk inhibitors *dusp2, dusp5* and *spry4* (*6*). Sproutys (Sprys) are known inhibitors of Ras/MAPKKK/MAPKK/ERK pathways and act by disrupting the complex of Growth-factor Receptor-Bound-2 protein (Grb2) and Sos and/or inhibiting the MAPKKK Raf (*23, 24*). Dusps are a subfamily of DUal-Specificity protein Phosphatases that regulate MAPKs negatively by dephosphorylating both tyrosine and threonine residues of the signature T-X-Y motif located within the kinase activation loop (*25*). Interestingly, inhibitors and their stability properties can impact the oscillation frequency of signaling pathways. For example, the frequency of the vertebrate segmentation clock is modulated by the stability of its core transcription factors together with Dusp inhibitors (*26, 27*). In scales, Erk inhibitors were predicted to control the maximum frequency of Erk waves by forming trailing regions of high-inhibitor concentration that are temporarily refractory to further Erk activation (*12*). Refractory behavior has been observed in many excitable systems, including firing neurons and muscles (*28, 29*). In this work, we set out to characterize if and how Erk modulate the transcripts of its own inhibitors to shape wave dynamics, frequency and ultimately tissue growth.

## Results and discussion

Simulations predict that refractory periods would exist because peaks of Erk activity are followed by peaks of Erk inhibitor that switch Erk off and prevent its further activation (*6, 12*). We decided to start the analysis of the relationship between Erk and its inhibitors by testing if wave behavior was compatible with the existence of a refractory period. A theoretical prediction deriving from the existence of a refractory period is that two travelling excitable waves would not cross each other when they meet, but merge instead (Fig. 1F Theory, Movie S4). To test whether Erk activity waves showed such a refractory behavior, we analyzed current and published datasets (*6*) searching for instances of colliding waves, and observed 9 such instances (Fig. 1F Experiment, Fig. S1C). In all these cases, waves did not cross each other but rather merged into an individual wave. These observations support the notion that Erk activity waves are followed by a high-inhibitor spatial refractory region, as predicted by theory (Fig. 1G). Theory predicts that the duration of the refractory period, and therefore the maximum wave frequency, depends on the effective lifetime of Erk inhibitors, which may include the transcription/translation times and RNA/protein lifetimes (Fig. 1G, H).

Theory predicts that refractory regions are formed by Erk stimulating its own inhibitors. We hypothesized that this modulation could take place at the transcript levels, given the long timescales involved (many hours; Fig. 1D, H; Fig. S1D), and that Erk activity waves would be followed by waves of inhibitor transcripts. To test this hypothesis, we set out to visualize the transcripts of the candidate Erk inhibitors *dusp2, dusp5, spry2* and *spry4* and relate them to Erk activity waves. To this end, we developed a single molecule Fluorescence In Situ Hybridization technique in intact regenerating scale osteoblasts (smFISH adapted from (*30*); Methods), achieving high signal-over-noise detection of transcripts in intact regenerating scale osteoblasts (Fig. 2A; Fig. S2A-D). This technique does not include any amplification step and allowed us to quantify the abundance of target transcripts directly by counting smFISH diffraction-limited spots (both nuclear and cytoplasmic) in z-maximum projections. We developed a pipeline to isolate the osteoblast tissue computationally, quantify smFISH spot density and relate it to Erk activity in the same scale (Fig. 2B).

**Figure 2.**
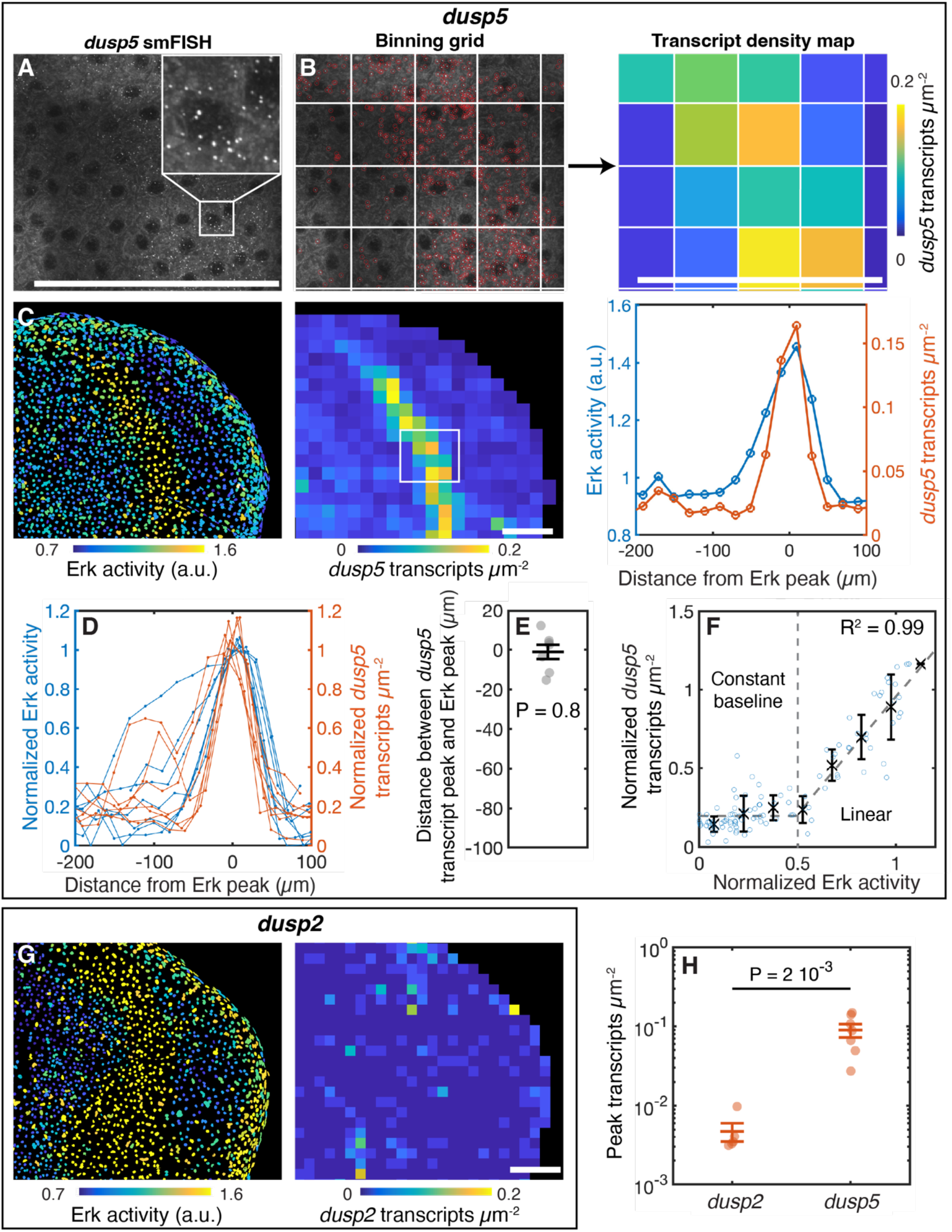
dusp5 transcripts are patterned in high density rings that co-localize with Erk activity waves, while dusp2 is expressed at lower levels. A. Example of a regenerating scale stained for dusp5 transcripts using smFISH (max projection), with magnification (inset). B. smFISH transcripts are automatically detected (left; red circles), binned (left; grid) and a transcript density map is calculated (right). C. Example of Erk activity and dusp5 transcript density in an individual scale (left: Erk activity map; middle: dusp5 transcript density map; right: spatial profiles of Erk activity and dusp5 transcript density). D. dusp5 transcript density and Erk activity profiles (n = 7 scales from 4 fish from 3 experiments). E. Distance between peaks of dusp5 transcript density and Erk activity profiles (with SEM; P-value of Student’s t-test against 0 µm is indicated; same dataset as in D). F. Quantification of the Erk activity-dusp5 transcript input-output relationship (with binned means (black crosses) and SD). Dashed line: ReLU fit (fitted parameters with 95% CI: C0 = 0.19 ± 0.02, δ = 1.6 ± 0.2, E0= 0.5 ± 0.1; the coefficient of determination R^2^ is indicated; same dataset as in D). G. Example of Erk activity (left) and dusp2 transcript density (right) in an individual scale. H. dusp2 and dusp5 peak transcript density levels (individual dots represent individual scales - dusp2: n = 5 scales from 3 fish from 2 experiments; dusp5: same dataset as in D); unpaired Student’s t-test P-value is indicated). The average peak value of dusp2 transcript density (5 10^-3^ transcripts/µm^2^) corresponds to ∼0.6 transcripts per cell. The average peak value of dusp5 transcript density (9 10^-2^ transcripts/µm^2^) corresponds to ∼13 transcripts per cell. a.u.: arbitrary unit. Scale bars, 100 µm.

We initially applied this technique to visualize *dusp5* transcripts (Fig. 2A, B, C). Strikingly, this analysis revealed that Erk activity waves co-localize with rings of enriched *dusp5* transcripts (Fig. 2C-E). This observation is compatible with a model in which Erk modulates either *dusp5* transcription or *dusp5* transcript lifetime, with a short *dusp5* transcript lifetime in either case. Indeed, since Erk activity waves move across scales at the roughly constant speed (∼20 µm h^-1^; Fig. S1D), we could convert spatial profiles into temporal dynamics using wave speed as conversion factor. As the standard deviation of the Erk activity-transcript peaks distance is ∼10 µm, we estimate transcript lifetime to be less than 30 min (Fig. 2E). Given the inferred short lifetime of *dusp5* transcripts with respect to Erk activity dynamics, the transcript production-degradation process can be considered at quasi-static equilibrium. Therefore, we computed the equilibrium input-output relationship between Erk activity and *dusp5* transcript density by comparing them in the same location. Remarkably, we observed a tight input-output relationship, with a characteristic Rectified Linear Unit response (ReLU, see e.g. (*31*); Fig. 2F). In a ReLU input-output relationship, the output stays at a constant baseline for inputs below a certain threshold, while it is linear above this threshold. In our measurements, transcript density was constant for roughly half of the Erk KTR sensor dynamic range, while it increases linearly for higher Erk KTR sensor values. This non-linear and threshold response is typical of processing units in neural networks which offers processing capabilities together with insensitivity to small perturbations (*31*).

This analysis suggests a ReLU response of *dusp5* transcript density to Erk activity. To test if *dusp5* transcript rings depended causally on Erk activity, we inhibited the upstream MAPKK Mek pharmacologically and quantified *dusp5* transcript profiles thereafter. Strikingly, *dusp5* rings resulted strongly impaired (Fig. S3A-E). Altogether, our results demonstrate that Erk activity waves induce rings of high levels of *dusp5* transcript.

After having characterized the relationship between Erk and *dusp5* transcripts, we set out to investigate the pattern of transcripts of the other *dusp* candidate, i.e. *dusp2*. As expected from previous bulk transcriptomics (*6*), *dusp2* transcripts, although detectable, exhibited strongly lower levels than *dusp5* and not clearly related to Erk activity waves (Fig. 2G, H; S4A-E). Given these results, we decided to not analyze *dusp2* transcripts further.

We continued our analysis of Erk inhibitors by analyzing Spry transcripts. In addition, to the bulk RNAseq candidate *spry4*, we analyzed the transcripts of *spry2*, which is expressed at similar levels as *spry4* in bulk RNAseq (*6*). As for *dusp5*, we discovered that Erk activity waves colocalized with rings of *spry2* higher transcript density (Fig. 3A-C), which indicated a short lifetime of s*pry2* transcripts (estimated <30min; Fig. 3C). An analysis of the Erk activity-*spry2* transcript simultaneous input-output relationship revealed a second instance of a ReLU response (Fig. 3D). While *dusp5* transcripts reached values an order of magnitude higher than the ReLU baseline (see Fig. 2F), *spry2* transcripts reached values just twice as high than the ReLU baseline (Fig. 3D). This suggests that Erk activity waves modulate *spry2* transcript density on top of a quantitatively comparable Erk-independent transcript baseline. As for *dusp5*, to test whether Erk regulates *spry2* transcripts, we inhibited Mek activity pharmacologically, findings that *spry2* peak levels were strongly inhibited (Fig. S5A-E).

**Figure 3.**
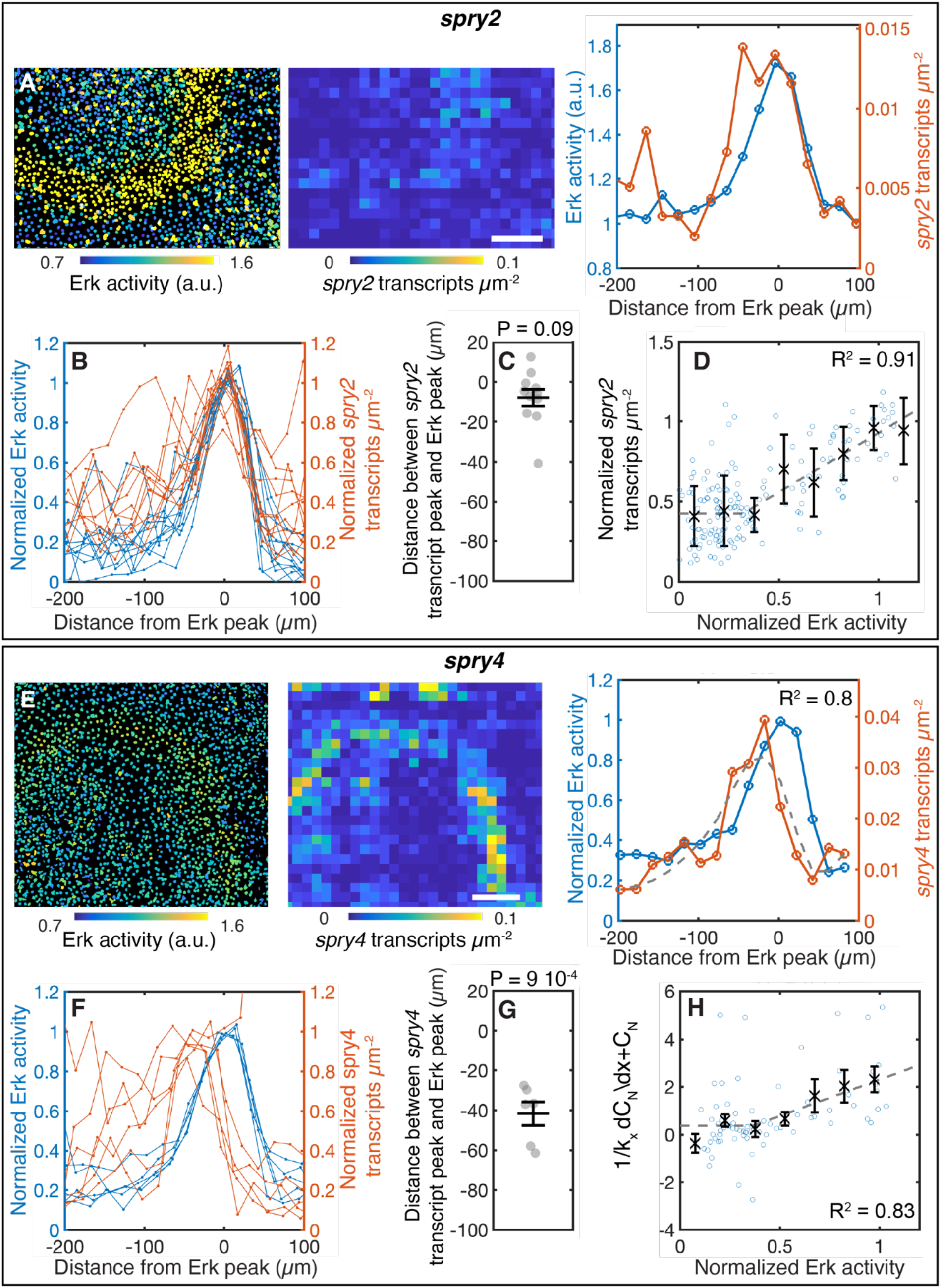
spry2 and spry4 transcripts are patterned in high density rings that, respectively, co-localize with and follow Erk activity waves. A. Example of Erk activity and spry2 transcript density in an individual scale (left: Erk activity map; middle: spry2 transcript density map; right: spatial profiles of Erk activity and spry2 transcript density). B. spry2 transcript density and Erk activity profiles (n = 11 scales from 5 fish from 4 experiments). The average peak value of spry2 transcript density (2 10^-2^ transcripts/µm^2^; see Fig. S6A) corresponds to ∼3 transcripts per cell. C. Distance between peaks of Erk activity and spry2 transcript density and Erk activity profiles (with SEM; P-value of Student’s t-test against 0 µm is indicated; same dataset as in B). D. Quantification of Erk-spry2 transcript input-output relationship (same dataset as in B, with binned means (black crosses) and SD). Dashed line: ReLU fit C = C0(1+ δ*H(E-E0)*(E-E0)) with 95% CI (fitted parameters: C0 = 0.4 ± 0.1, δ = 2 ±1, E0= 0.2 ± 0.1; the coefficient of determination R^2^ is indicated). E. Example of Erk activity and spry4 transcript density in an individual scale (left: Erk activity; middle: spry4 transcript density map; right: normalized Erk activity and spry4 transcript density spatial profiles. Dashed line: fit with an ODE model in which Erk modulates spry4 transcript production (see Methods; the coefficient of determination R^2^ is indicated). F. spry4 transcript density and Erk activity profiles (n = 6 scales from 5 fish from n = 4 experiments). The average peak value of spry4 transcript density (4 10^-3^ transcripts/µm^2^; see Fig. S6A) corresponds to ∼6 transcripts per cell. G. Distance between peaks of spry4 transcript density and Erk activity profiles (with P-value of Student’s t-test against 0 µm; with SEM, same dataset as in F). H. Quantification of Erk-spry4 transcript input-output relationship (with binned means (black crosses) and SD). Dashed line: fitted Erk-spry4 transcript input-output relationship with ODE model in which Erk modulates spry4 transcript production (see Methods; fitted parameters are shown in Fig. S6B, C; the coefficient of determination R^2^ is indicated; same dataset as in F). a.u.: arbitrary unit. Scale bars, 100 µm.

We then turned to likewise analyzing *spry4* transcripts, which also correlated with Erk activity waves and exhibited similar levels than *spry2* (Fig. 3E, F; S6A). However, in contrast to *dusp5* and *spry2* transcripts, *spry4* transcript density peaks follow Erk activity waves with a spatial delay of roughly 40 µm, corresponding to a temporal delay of roughly 2h (Fig. 3G). The distance between peaks in Erk activity and *spry4* transcript density indicates that the lifetime of *spry4* transcripts is not negligible with respect to Erk dynamics. Thus, to calculate the input-output relationship between Erk and *spry4* transcripts, we needed to fit the *spry4* transcript spatial profile with an ordinary differential equation model which included transcript production and degradation. In the model, transcript production was hypothesized to depend on the measured Erk activity in the same location, through a ReLU response (Methods). This model could fit the *spry4* profiles (Fig. S6B, C). The inferred input-output relationship between Erk activity and *spry4* transcription revealed a low baseline of *spry4* transcription with respect to peak (Fig. 3H). Of note, *spry4* transcript profiles could also be fitted with an alternative model in which Erk modulated *spry4* transcript lifetime instead of transcription (Methods). Finally, as for *dusp5* and *spry2*, we analyzed *spry4* transcript profile following pharmacological Mek inhibition. We discovered that Mek inhibition strongly inhibited *spry4* transcript rings (Fig. S6D-H). Altogether, these results demonstrate that Erk activity waves modulate their own inhibitors *dusp5, spry2* and *spry4*, which form trailing waves of high transcript density. In addition, we discovered non-linear Rectified Linear Unit input-output responses in these relationships.

These results demonstrated the transcriptional effect of Erk activity waves on their own inhibitors and thus on their own excitability machinery. In many systems, Erk activation can modulate several other cellular processes, such as a cell differentiation (*32, 33*). In scales, osteoblasts grow and mature as scales regenerate, which is favored by the master regulator transcription factor *osterix*, which is an Erk target in other systems (*34-36*). Thus, we asked whether Erk activity waves had also effects on osteoblasts maturation through modulating *osterix* transcription. We reasoned that this would offer an intriguing mechanism to couple scale growth, which is induced by Erk activity waves (*6*), with osteoblast maturation throughout regeneration. To test this hypothesis, we turned to analyzing the pattern of *osterix* transcripts with smFISH. As expected, we detected a high number of *osterix* transcripts in the osteoblast tissue (Fig. 4A). In nuclei, we often observed one or two large spots that we interpreted as active transcription sites (Fig. 4A, B top; arrowheads). Strikingly, quantification of the number of active transcription sites per cell revealed rings of higher active transcription following Erk activity waves at about 50 µm distance (Fig. 4B-E), indicating the modulation of *osterix* transcription. Compatibly with this observation, we discovered rings of high *osterix* transcript density following Erk activity waves (Fig. 4C, F, G). As done previously, we tested a causal role of Erk activity in the modulation of *osterix* transcription and transcript density by analyzing their patterns following Mek inhibition (Fig. S7A). We discovered that rings of *osterix* active transcription and transcript high density were strongly inhibited following treatment with a Mek inhibitor (Fig. S7B, C). Interestingly, after Mek inhibition, *osterix* active transcription site and transcript density showed a graded profile, high at the scale boundary and decreasing toward the interior of the scale (Fig. S7B right, S7C right). Overall, these experiments showed that Erk activity waves modulate *osterix* transcription over a graded Erk-independent baseline of transcriptional activity.

**Figure 4.**
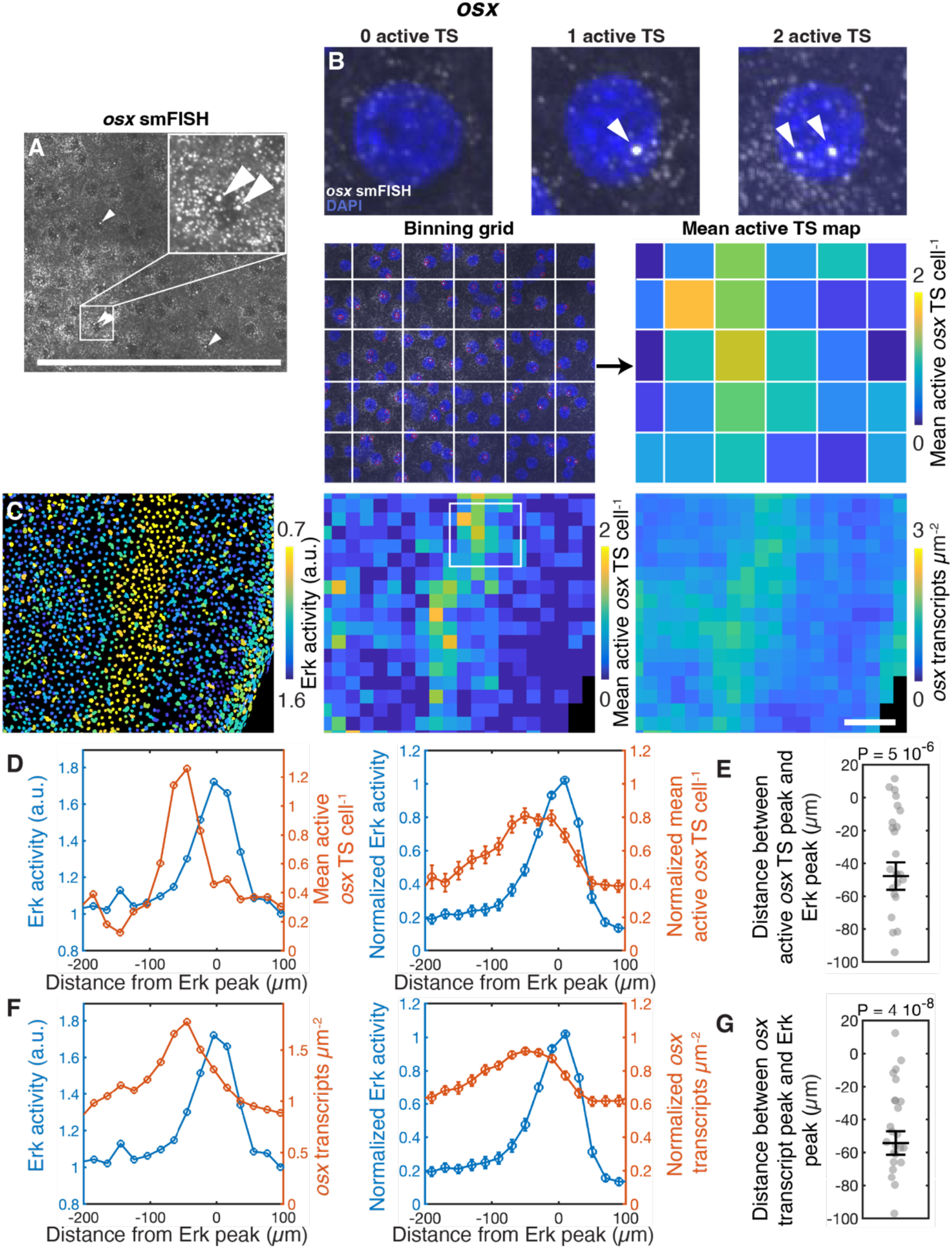
osx transcripts and active transcription sites are patterned in high density rings that follow Erk activity waves with a delay. A. Example of a regenerating scale stained for osx transcripts using smFISH (max projection) with magnification (inset; white arrows: examples of active transcription sites (TS)). B. Examples of osteoblasts scale co-stained with DAPI (blue) and osx smFISH (white) (arrowheads: TS). A binning grid (bottom left; red circles: detected TS) is used to calculate a map of the mean number of osx active TS per cell (bottom right; thereafter: TS per cell). C. Example of Erk activity, osx TS per cell and osx transcript density in an individual scale (left: Erk activity map; middle: map of osx TS per cell; right: osx transcript density map). D. Example of spatial profile of osx TS per cell with Erk activity in an individual scale (left) and average of multiple scales (right; with SEM; n = 28 scales from 15 fish; n = 13 experiments). E. Distance between peaks in osx TS per cell and Erk activity (with SEM; the P-value of Student’s t-test against 0 µm is indicated; same dataset as in D). F. Example of quantification of osx transcript density with Erk activity in an individual scale (left; same dataset as in D) and average of multiple scales (right; with SEM; same dataset as in D). The average peak value of osx (1.7 transcripts/µm^2^) corresponds to ∼250 transcripts per cell. G. Distance between peaks in Erk activity and osx transcript density (with SEM; the P-value of the Student’s t-test against 0 µm is indicated; same dataset as in D). a.u.: arbitrary unit. Scale bars, 100 µm.

Overall, these results revealed that Erk activity waves modulate their own inhibitors through transcript pattern modulation. This result is potentially in opposition with other cases of faster Erk activity waves (*4, 5, 8-10*), in which likely Erk activity would not be sustained long enough to allow for the activation of transcription. Our findings show that such a modulation occurs through a non-linear ReLU function, which is typical of processing units in neural networks. This constant-then-linear response can provide high linear-like sensitivity, while not respond to small perturbations. Molecularly, this type of response could emerge from Erk being sequestered by a biding site that would be alternative to the Erk targets that activate inhibitor transcription; in this model, Erk would activate transcription of its inhibitors, linearly, when the concentration of active Erk overcomes the abundance of Erk “sequestering sites”. It is important to note that, in the case of Erk inhibitors, our observations are also compatible with Erk modulating the pattern of transcripts by regulating their lifetime, instead of transcription rate; this type of transcript lifetime modulation was observed, for example, in melanoma and Hela cells (*37, 38*).

Strikingly, we discovered that Erk activity waves modulate their own maturation via inducing transcription of the transcription factor *osterix*. This finding suggests a model in which osteoblast maturation could act as a “counter” of Erk activity waves; in this model, the more waves travel across the tissue, the more osteoblasts mature through *osterix* expression. Such osteoblast maturation may further modulate the expression and transcript lifetimes of Erk inhibitors. In turn, this modulation could limit the maximum frequency of Erk activity waves. Overall, this mechanism could tune the frequency of Erk activity waves across regeneration and eventually terminate their propagation.

Altogether, through spatially and temporally resolved experiments and quantifications, we demonstrated that Erk activity waves have transcriptional effects on their own inhibitory machinery and on osteoblast maturation. These results demonstrate that Erk activity waves are self-regulating and have impact on the bone morphogenetic program in a regenerating vertebrate appendage. We expect that similar principles may operate in different Erk activation patterns, timescales and contexts.

## Materials and Methods

### Fish husbandry, scale injury and pharmacological treatments

Zebrafish deriving from Ekkwill, Ekkwill/AB and AB strains were maintained between 26 and 28.5 °C with a 14:10 h light:dark cycle. Fish between 3 and 18 months old were used for experiments. Scale plucking was performed essentially as previously described (*6, 15, 39*). In brief, fish were anaesthetized in 0.09% phenoxyethanol (Sigma L77699) or 0.032% Tricaine (Sigma-Aldrich E10521) in system water until swimming stopped and operculum movement slowed. Then, they were placed in a Petri dish, and fluorescent scales were viewed with a fluorescence dissecting microscope. Three rows of 7–15 scales were plucked with forceps from the trunk of the fish. After scale removal, fish were returned to system water to recover from anesthesia. No statistical methods were used to predetermine sample size. Males and females were randomly allocated in control and experimental groups. When possible, siblings were used for control and experimental groups. It was not possible to blind investigators during data collection, because fish from control and experimental groups can be distinguished by cell behaviors and/or fluorescent reporters. However, data quantification was mainly performed automatically using the same computational algorithm. When necessary, human manual data curation was performed by blinded researchers, although often data from control and experimental groups can be recognized from phenotypes. Researchers were not blinded during data visualization. Fish husbandry and animal experiments were approved by the Service de la Consommation et des Affaires Vétérinaires of the République et Canton de Genève (GEH8A and GE225A-H/GE547, respectively). Transgenic lines used in this study were: (*osx:H2A-mCherry*)^pd310^ (*15*) and Tg(*osx:ErkKTR-mCerulean*)^pd2001^ (*6*). Re-analyzed datasets from (*6*) included the transgenes: Tg(*osx:mCherry-zCdt1*)^p270^ (*15*), Tg(*osx:Venus-hGeminin*)^p271^ (*15*), Tg(*osx:H2A-mCherry*)^pd310^ (*15*) and Tg(*osx:ErkKTR-mCerulean*)^pd2001^ (*6*).

For pharmacological treatments by immersion, fish were immersed in the pharmacological compound diluted to working concentration in fish water for the duration of the treatment. In each experiment, control fish were treated with the same concentration of DMSO vehicle as the treated group and collected on the same day. In the case of PD0325901 (Selleck Chemicals S1036) (stock concentration: 100 µM; working solution: 10 µM) and DMSO control, fish were immersed for 9h in the treatment solution. Scales with Erk activity waves at a similar distance from the scale border were selected for the DMSO- and PD0325901-treated groups. During treatment, fish were maintained off the aquarium system in the dark, kept in 0.7l tanks at a density of 2 fish per tank. Thereafter, fish were euthanized, scales collected and smFISH detection and imaging were performed as described below.

### Erk activity imaging and quantification

In vivo confocal imaging was performed essentially as previously described (*6, 15, 39*). In brief, a zebrafish was anaesthetized in 0.016% tricaine in system water and transferred to a 1% agarose bed in 3D-printed black ABS (Acrylonitrile Butadiene Styrene) plate. The caudal fin of the fish was set on a glass slide or a ABS piece to bring the caudal peduncle of the trunk parallel with the platform, and diluted tricaine was placed near the head of the fish. Cooling 1% agarose was applied on the caudal fin, the trunk anterior to the imaged scale, and the areas of platform dorsal and ventral to the scale. Then, fish were immersed in diluted tricaine. Gill movements were monitored visually and, when they slowed, system water was applied using a peristaltic pump (CONTROL COMPANY Chemical Metering Pump; Model 3385; silicon tubing: Tygon, 2.4 mm inner diameter and 4mm outer diameter, E-3603) until regular rhythm was restored. In each fish, the same scale was recognized at different time points from its position in the scale array and/or morphology. After imaging, a system water flow was applied using the peristaltic pump, until gill movement quickened. Then, the fish was returned to system water. For longitudinal time courses, fish were mounted, imaged and then returned to system water at each time point. For *in vivo* or fixed scales imaging, confocal images were acquired using a Leica Stellaris 5 confocal microscope. As scales are larger than the field-of-view of our microscopy setup, they were imaged with multiple overlapping z-stacks (1–40 z-stacks with variable number of planes) to cover the entire osteoblast tissue. Fluorescent proteins were imaged using the following lasers: Erk KTR–mCerulean, 448 nm; H2A–mCherry, 561-580 nm (White Light Laser). A variable laser power and gain was used depending on the expression of the transgenic reporters. DAPI was imaged using a 405 nm laser. Shown raw images were adjusted to enhance contrast.

Erk quantification from confocal images was performed as in (*6, 39*) using custom-written MATLAB (Mathworks) 2024b software. In brief, scale stacks were stitched, rotated in 3D to align them to the x-y plane and several overlapping neighboring scales were computationally removed using a semi-automated routine. Therefore, several image processing steps are required to isolate the dermal osteoblast monolayer that is the subject of this study. Thereafter, nuclei were segmented from *osx*:H2A-mCherry images using a custom-written pipeline and the gaussian-mixture model TGMM (*40*). A mask corresponding to each nucleus was drawn using nuclei segmentation; the nuclear mask was dilated, and the difference of the dilated mask and the nuclear mask was taken as a cytoplasmic mask. Then, average *osx*:ErkKTR–mCerulean fluorescence was calculated in the nuclear and cytoplasmic regions. Erk activity was measured as the ratio of cytoplasmic and nuclear average Erk KTR–mCerulean fluorescence levels. Wave speed was calculated as in (*6*). The initial distance of Erk activity waves to the scale rim before treatment with DMSO or PD0325901 in Mek inhibition experiments was measured manually using FIJI (*41*).

All untreated fish were imaged 4 days post-plucking. In the case of pharmacological treatments, treatment started 4 days post-plucking and fish were imaged 5 days post-plucking; then fish were euthanized, scales collected and the smFISH protocol was performed.

### smFISH

Single molecule RNA Fluorescence In situ Hybridization was performed adapting a protocol from (*30*) to achieve high signal-over-noise detection of RNA transcripts in intact tissue. Probes were designed using eFISHent software (https://github.com/BBQuercus/eFISHent) and ordered from Microsynth DNA synthesis.

Thereafter, DNA oligos were labeled with fluorescent dyes using Terminal Deoxynucleotidyl Transferase enzyme. Probes were designed to recognize exonic regions of *spry2 spry4, dusp2, dusp5* and *osx* transcripts.

After euthanasia, scales were collected in 1X PBS and fixed in a 4% paraformaldehyde solution (EMS 15735-10S) in 1X PBS. Then, scales were permeabilized in 0.5% Triton-X (Fisher bioreagent BP151-100) in 1X PBS. For smFISH detection, scales were placed on coverslips. Scales were equilibrated in washing buffer. Scales were treated in a hybridization solution, then washed with washing buffer, stained with DAPI solution (Sigma-Aldrich D9542-10MG; 1 µg/ml in 1x PBS) for 20 min and mounted in Prolong Gold Antifade Reagent (Cell Signaling Biotechnology 9071S).

### smFISH. Imaging, data curation and spot detection

First, DAPI and *osx*:ErkKTR-mCerulean signal were imaged on a Stellaris 5 confocal microscope as described in “Erk activity imaging and quantification”. Thereafter, the smFISH probes and DAPI signal were imaged on either of the following spinning disk microscopes: a Zeiss Axio Observer 7 equipped with a CSU-W1 automatic Yokogawa spinning disk head); or an inverted Nikon Ti with a CSU-W1 Yokogawa spinning disk head. 3D stacks were typically acquired with adapted exposure times and laser power. Shown raw smFISH images were adjusted to enhance contrast.

smFISH quantifications were performed on z-maximum projections of curated stacks. Stacks were curated computationally using a custom-written FIJI macro to isolate the osteoblast tissue that is the subject of this study (hyposquamal; dermal side) from the episquamal layer (epidermal side) and other neighboring tissues. To this end, scale anatomy and tissue layers could be identified using the smFISH probe signal (spots and background signal). Of note, during this process, high intensities cell debris that would generate false positive detected spots were also removed. After data curation, the resulting segmentation mask was correspondingly upsized and applied to the original stacks.

smFISH spots were detected from smFISH images in z-maximum projections using the FIJI plugin TrackMate (*42*). smFISH spots were interpreted as individual transcripts or transcription sites based on their size. Detection thresholds were custom set visually. In a subset of samples, detection thresholds were selected by two researchers independently, obtaining similar results.

### smFISH. Transcript/transcription site profile quantification and comparison with Erk activity profiles

Transcript density maps/profiles and active transcription sites per cell maps/profiles, as well as Erk activity profiles, were obtained using a custom-written MATLAB pipeline (version 2024b). smFISH and Erk activity images were obtained on different microscopes (see “*smFISH. Imaging, data curation and spot detection”*). Therefore, smFISH and Erk activity images were registered automatically or semi-automatically using nuclear DAPI signal, which was imaged on both microscopes, as a reference. Erk activity was quantified for individual cells as in *“Erk activity imaging and quantification”*.

Transcript density maps were computed from the spatial coordinates of individual transcripts detected by smFISH. Transcript density was obtained by binning transcripts in 2D and dividing by bin area. The maps of average number of transcripts/active transcription sites per cell were computed by assigning each transcription site to the nearest nucleus centroid, then binning nuclei in 2D and averaging their number of assigned transcripts/transcription sites.

Transcript density linear profiles were calculated as follows. A ROI for the scale was manually selected and, when necessary, further restricted to exclude regions in which the tissue was damaged by the staining procedure, showing excess debris or the image was not complete or damaged. The ROI was restricted to the region in which both the Erk KTR and smFISH signals were imaged. An Erk activity wavefront was manually selected as a segmented line based on the Erk activity quantification. We restricted the analysis to travelling waves away from the source. Transcript density linear profiles were quantified by computing the shortest distance of each transcript to the wavefront using a custom function. Transcripts whose minimum-distance coordinate was on the edge of the wavefront were classified as outside the front and excluded. Remaining transcripts were binned by their distance to the wavefront and a profile of transcripts density as a function of distance from the wavefront was calculated.

When calculating the profiles of average number of transcripts/active transcription sites per cell, the distance between each nucleus centroid and the wavefront was calculated; nuclei were binned depending on this distance and the number of transcripts/active transcription sites per cell of nuclei in each bin was averaged. A similar procedure was performed to calculate Erk activity profiles, in this case by averaging Erk activity values assigned to each nucleus.. The same procedures were used to calculate the profiles of number of transcripts/active transcription sites/Erk activity away from the scale edges. In this case, the wavefront was substituted with a segmented line following the scale edge. For all transcript density profiles, points were excluded when less than 5 transcripts per bin were counted. For profiles of number of transcripts/active transcription sites per cell and Erk activity, points were excluded when less than 5 nuclei per bin were counted.

Profile peaks were calculated using parabolic fits on manually selected regions and a Savitzky-Golay smoothening. If reliable fitting was not possible, peaks were manually selected. The normalization of profiles of transcript density and number of active transcription sites per cell was performed by dividing by the fitted profile peak. Erk activity profile normalization was performed by clamping between the minimum value of the profile and the fitted peak. Peak number of transcripts per cell values were calculated from peak transcripts per μm^2^ values, using a conversion factor calculated as the ratio between the ROI area and the number of cells in the ROI, averaged for scales of each probe.

### Fit of transcript density profiles

The smFISH transcript density profile of each scale was fitted with the solution of an ordinary differential equation (ODE):

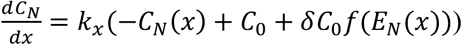

where *C*_*N*_(*x*) and *E*_*N*_(*x*) are the normalized spatial profiles of transcript density and Erk activity (binned averages; see above for normalization). The parameters are: *k*_*x*_ that is the transcript spatial lifetime, equal to the transcript lifetime multiplied by wave velocity; *C*_0_ that controls the baseline transcript production rate *k*_*x*_*C*_0_; *δ* that controls the coefficient *k*_*x*_*δC*_0_ of the Erk-dependent ReLU transcript production rate *f*(*E*_*N*_(*x*)) = *H*(*E* − *E*_0_)(*E* − *E*_0_) where H is the Heaviside function. When the ODE is in a quasi-static equilibrium, it reduces to:

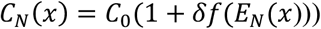

which was used to fit the Erk-transcript density input-output relationship for *spry2* and *dusp5*, since transcript density and Erk activity profile peaks colocalized, as expected in a quasi-static regime. In the case of *spry4*, the transcript density peak followed that of the Erk activity profile and thus the ODE could not be taken at a quasi-static equilibrium. Therefore, we fitted the *C*_*N*_(*x*) and *E*_*N*_(*x*) couples using the full ODE. The fit was performed using a custom-written MATLAB code based on the ODE solver ode45 and minimization of a quadratic error function using the function fmincon. The coefficient of determination is calculated as 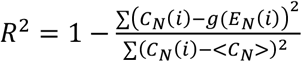 where *g*(*E*_*N*_ (*i*)) is the solution of the ODE equation.

Curves could also be fitted with a model in which Erk modulates transcript lifetime:

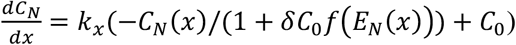

with parameters *k*_*x*_, *C*_0_ and *δ*. This second model gives the same quasi-static equilibrium relationship as in the model in which Erk modulates the transcript production rate.

### Mathematical model of Erk activity waves excitability

The mathematical model of Erk activity waves excitability is based on the reaction-diffusion model presented in (*6, 12*) and solved numerically using a forward Euler method as described. The mathematical model includes three variables: non-diffusible Erk activity E, a diffusible Erk activator A and a non-diffusible Erk inhibitor I. The equations used in this work are:

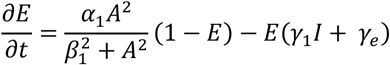

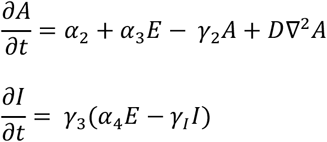

where *α*_1_ is the Erk activation rate at saturating activator, *β*_1_ is the AC_50_ of Erk activation by the activator, *γ*_1_ is the Erk inactivation rate by the inhibitor I, *γ*_*e*_ is the Erk autonomous de-activation rate, *α*_2_ is the activator production rate (positive in the source region, null outside), *α*_3_ is the Erk-dependent activator production rate, *γ*_2_ is the activator autonomous degradation rate, *γ*_3_*α*_4_ is the inhibitor production rate and *γ*_3_*γ*_*I*_ is the inhibitor autonomous degradation rate. The parameters used are: *α*_1_: 12.6 h^-1^, *β*_1_: 0.35, *γ*_1_: 15.4 h^-1^, *γ*_*e*_: 0.14 h^-1^, *α*_2_: 0.112 h^-1^ in the source region (null outside), *α*_3_: 3.9 h^-1^, *γ*_2_: 11.8 h^-1^, *D*: 566 µm^2^/h, *γ*_3_: 0.14 h^-1^, *γ*_I_: 0.4 h^-1^ or 1 h^-1^ and *α*_4_: 0.5. The simulation domain is a 1090 × 1090 μm square (simulation grid size of 5 μm; time-step of 0.01s) in the standard, spiral and colliding waves simulation (Fig. 1D-F). For the parameter sensitivity analysis of Fig. 1H, a simulation grid size of 15 μm and time-step of 0.01 s were used. The scale region is a disc that is 520 μm in radius. Erk activity and inhibitor concentrations are null outside the scale region. The activator diffuses outside the scale, but absorbing boundary conditions are set at the domain boundary.

### Statistical Analysis

When relevant, the statistical test used, number of samples and independent experiments is specified in the corresponding figure legends.

## Acknowledgments

The authors thank Pierre Gönczy, Konrad Marx and Ivan Rodriguez for critical reading of the manuscript and/or discussion.

## Author contributions

Conceptualization: CC, TG, ADS

Methodology: CC, TG, FV, ADS

Software: TG, ADS

Data curation: CC, TG, ADS

Validation: CC, TG

Formal analysis: CC, TG, ADS

Investigation: CC, TG, ADS

Visualization: TG, ADS

Supervision: ADS

Writing—original draft: CC, TG, ADS

Writing—review & editing: CC, TG, FV, ADS

Projects administration: ADS

Funding acquisition: FV, ADS

