## Supplementary Information for "Transcriptional feedback of Erk signaling waves in zebrafish scale regeneration"

### Table of contents

**Supplementary Movies Legends** …………………………………………………………………………. **2**

**Supplementary Figures** …………………………………….…………………………………….…. **3**

#### Supplementary Movies Legends

**Movie S1**
Example of an Erk activity wave in theoretical simulations. Erk activity and inhibitor concentration are shown; an Erk activity wave is followed by a wave of inhibitor. Corresponds to Fig. 1D (Theory). Scale bar: 200 µm.

**Movie S2**
Example of a spiral Erk activity wave in theoretical simulations. Erk activity and inhibitor concentration are shown; a spiral Erk activity wave is followed by a spiral wave of inhibitor. Corresponds to Fig. 1E (Theory). Scale bar: 200 µm.

**Movie S3**
Example of a spiral Erk activity wave in vivo. Corresponds to Fig. 1E (Experiments). Scale bar: 200 µm.

**Movie S4**
Example of two colliding Erk activity waves in theoretical simulations. Erk activity and inhibitor concentration are shown. Two Erk activity waves collide and merge. (left). Corresponds to Fig. 1F (Theory). Scale bar: 200 µm.

#### Supplementary Figures

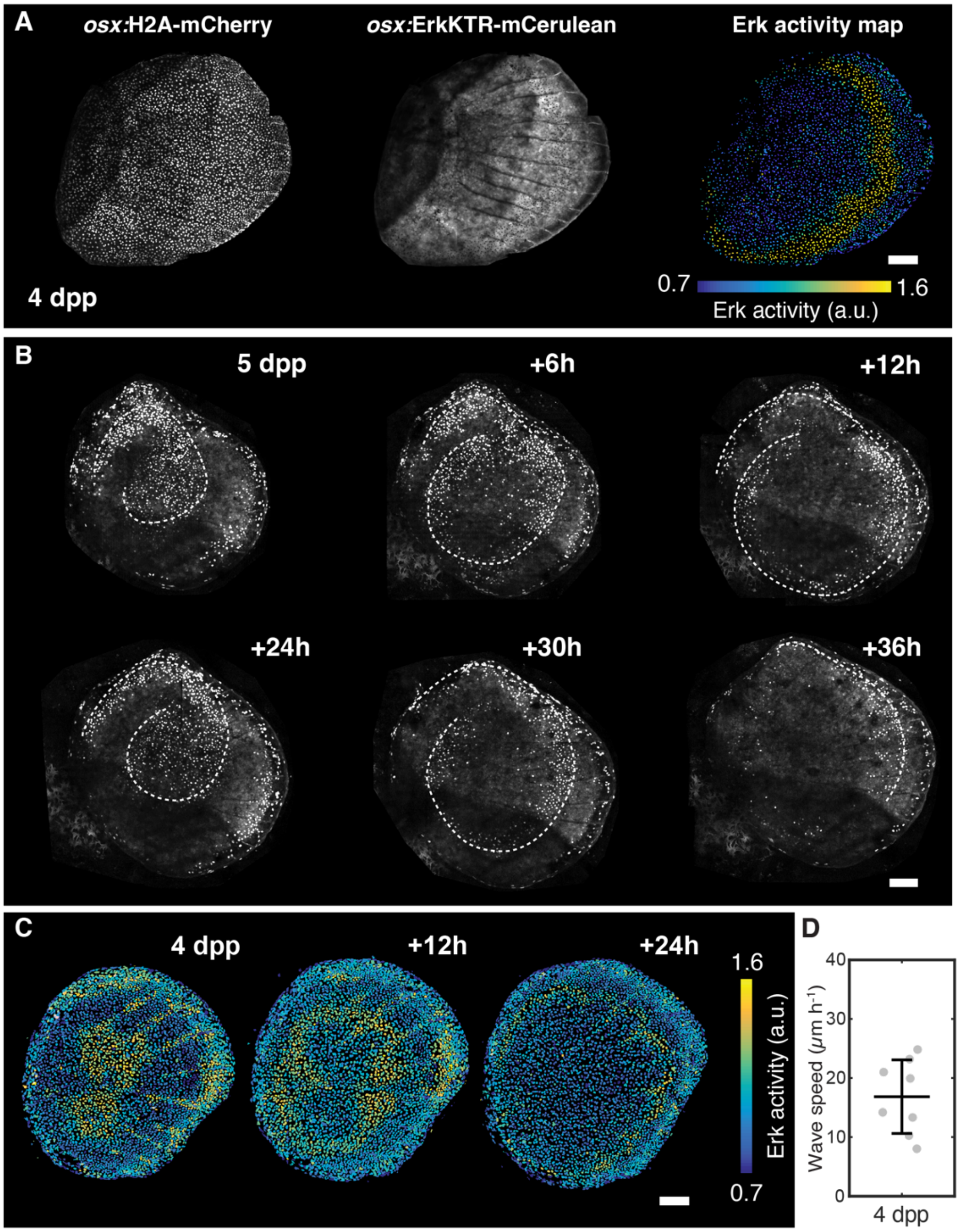


***Figure S1 – Measurement of Erk activity in regenerating zebrafish scales. Additional examples of spiral and colliding Erk activity waves. Speed of Erk activity waves.***

*A. Example of quantification of Erk activity in scales of fish expressing the live biosensor Erk KTR (nuclear marker* osx:*H2A-mCherry (left); Erk biosensor* osx:*ErkKTR-mCerulean (middle); Erk activity map (right)). B. Example of two subsequent spiral waves in the signal of* osx:*Venus-hGeminin (from a re-analyzed dataset from (1)).* osx:*Venus-hGeminin is a component of the cell-cycle reporter FUCCI (7, 31) and follows Erk activity waves (1). C. Additional example of two Erk activity waves colliding and merging (see Fig. 1F; this data was partially re-analyzed from (1)). D. Erk activity waves speed at 4 days post-plucking. Average speed: (17 ± 6) µm/h (n = 8 scales from 7 fish from two experiments). a.u.: arbitrary unit. dpp: days post-plucking. Scale bars, 100 µm.*

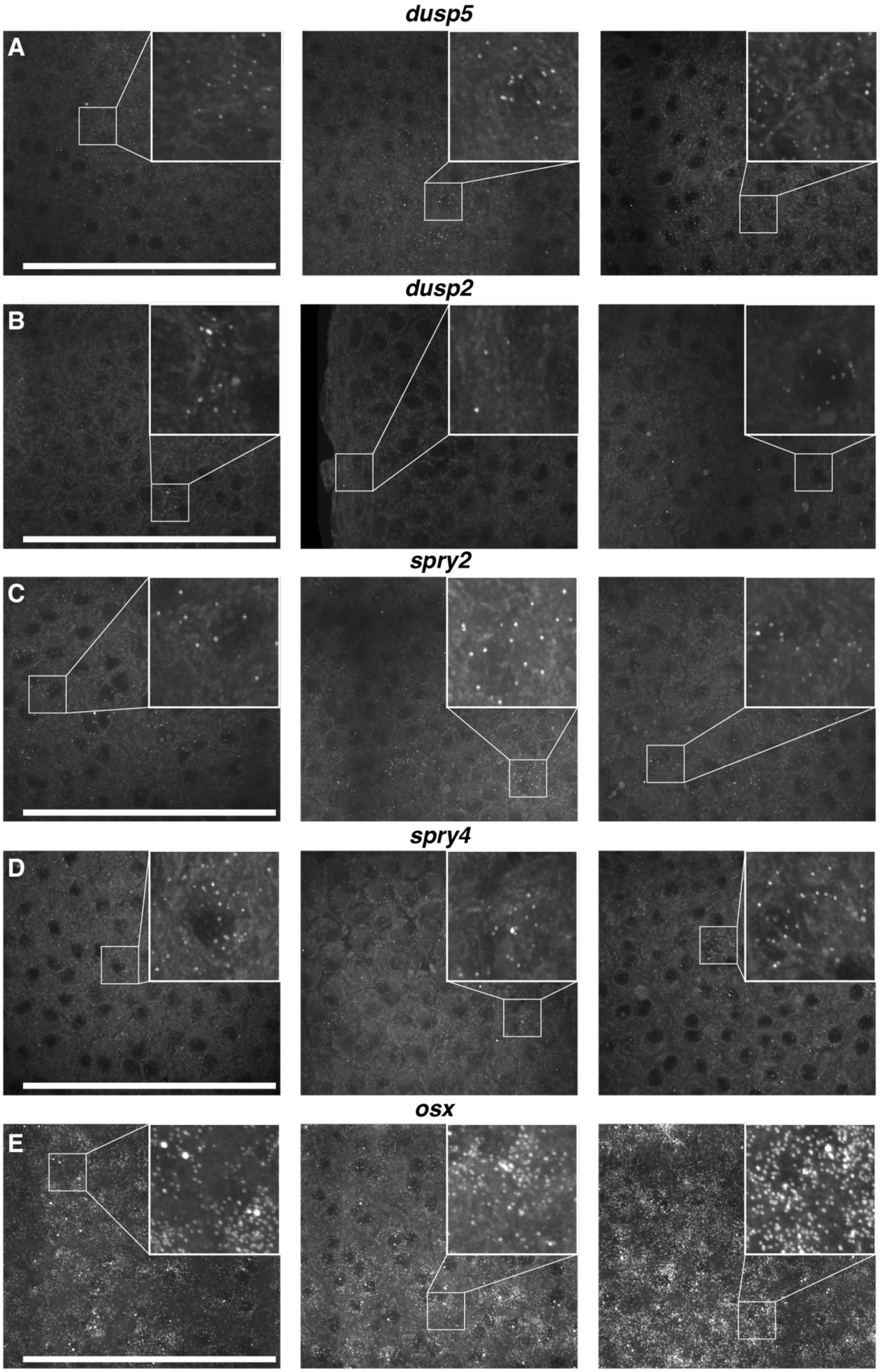


***Figure S2 – Examples of smFISH transcript detection in regenerating scales.***
*A-E. Examples of smFISH for* dusp5*,* dusp2*,* spry2*,* spry4 *and* osx *transcripts. Insets: magnification. Max projections are shown. Scale bars, 100 µm.*

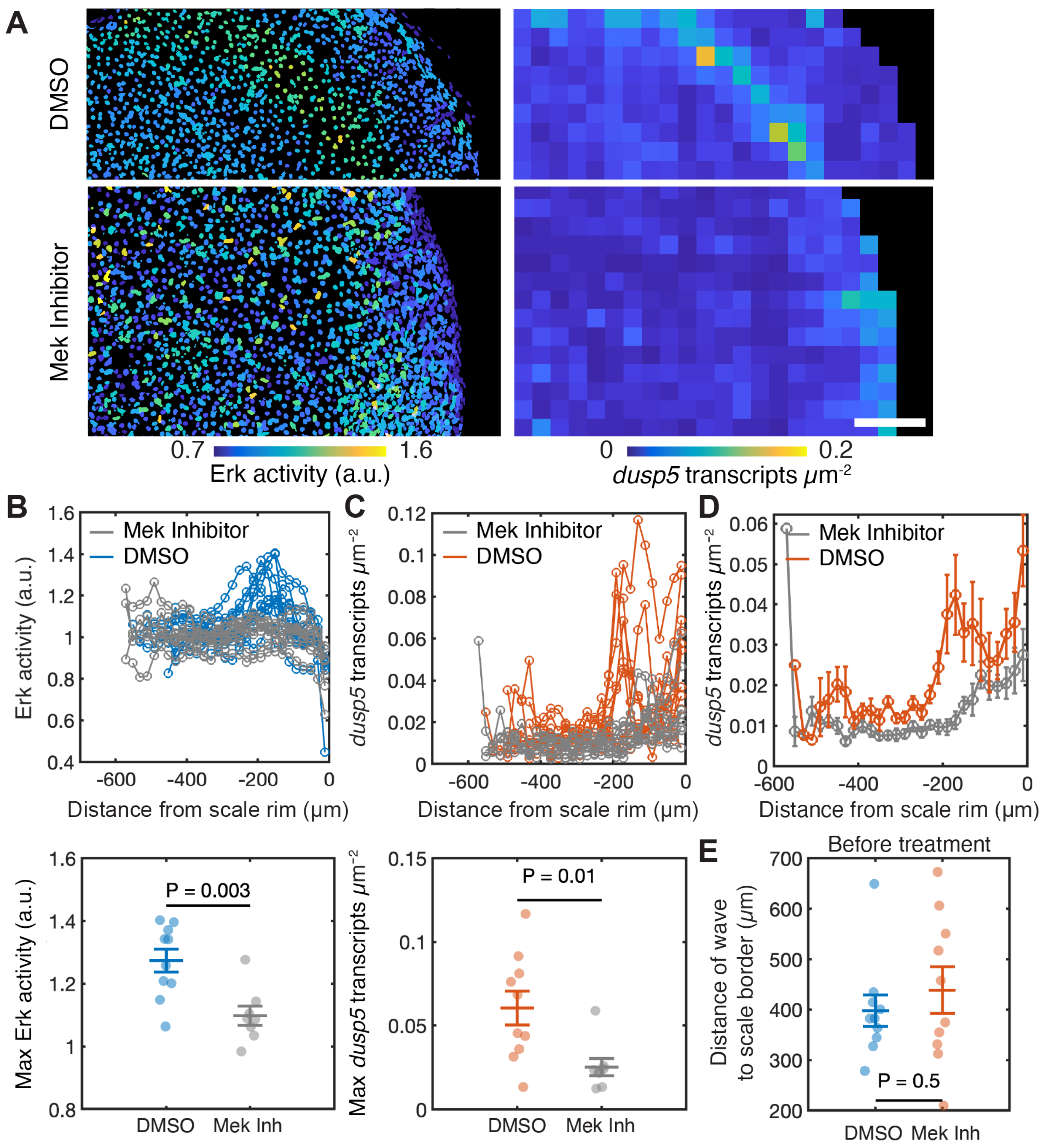


***Figure S3 –* dusp5 *transcript rings are impaired in scales treated with a Mek inhibitor.***

*A. Example of Erk activity and* dusp5 *transcript density in control scales (treated with DMSO) and scales treated with the Mek inhibitor PD0325901 (10 µM). B, C. Erk activity (B, top) and* dusp5 *transcript density (C, top) in control scales (treated with DMSO) and scales treated with the Mek inhibitor PD0325901 (10 µM). Amplitude of peaks of Erk activity (B, bottom) and* dusp5 *transcript density (C, bottom) (with unpaired Student's t-test P-value). DMSO: n = 10 individual scales from 2 fish from a single experiment; Mek inhibitor, n = 8 individual scales from 2 fish from a single experiment. D. Average* dusp5 *transcript density in control scales (treated with DMSO, same dataset as in B, C) and scales treated with the Mek inhibitor PD0325901 (10 µM; same dataset as in B, C). E. Distance of the wavefront to the scale border before treatment with DMSO or with the Mek inhibitor PD0325901 (with unpaired Student's t-test P-value treated with DMSO, same dataset as in B, C). a.u: arbitrary unit. Scale bar, 100 µm.*

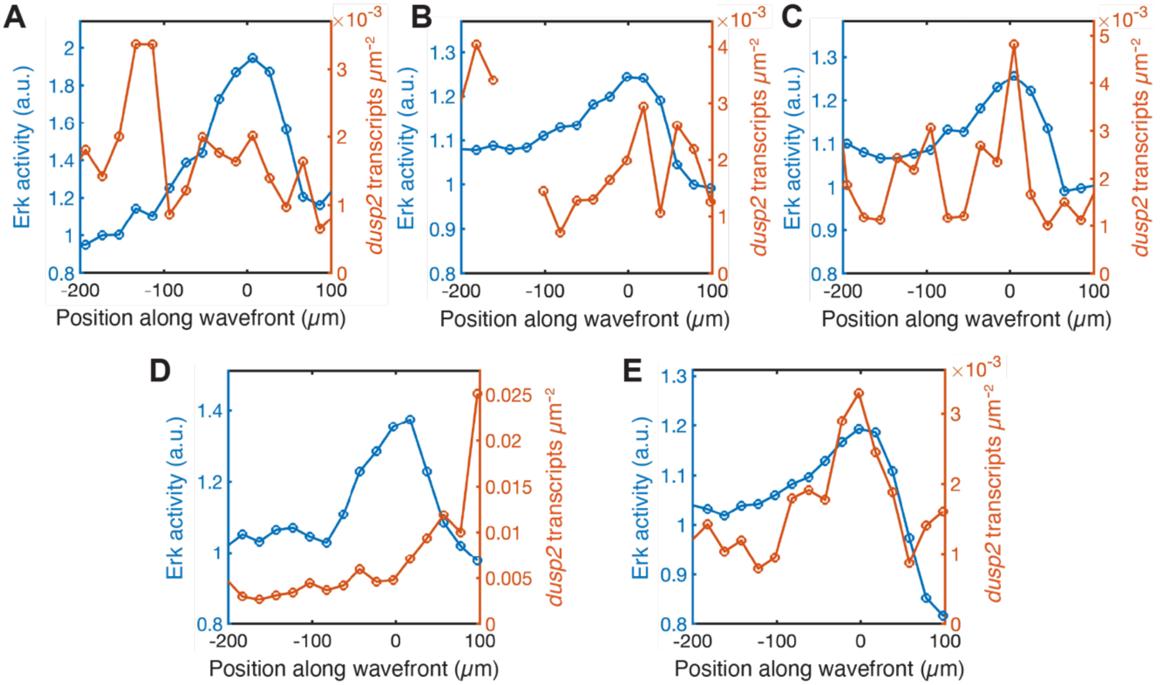


***Figure S4 – Examples of Erk and* dusp2 *transcript density spatial profiles.***

*A. Erk and* dusp2 *transcript density spatial profiles (example shown in Fig. 2G). B-E. Examples of Erk and* dusp2 *transcript density spatial profiles. a.u: arbitrary unit.*

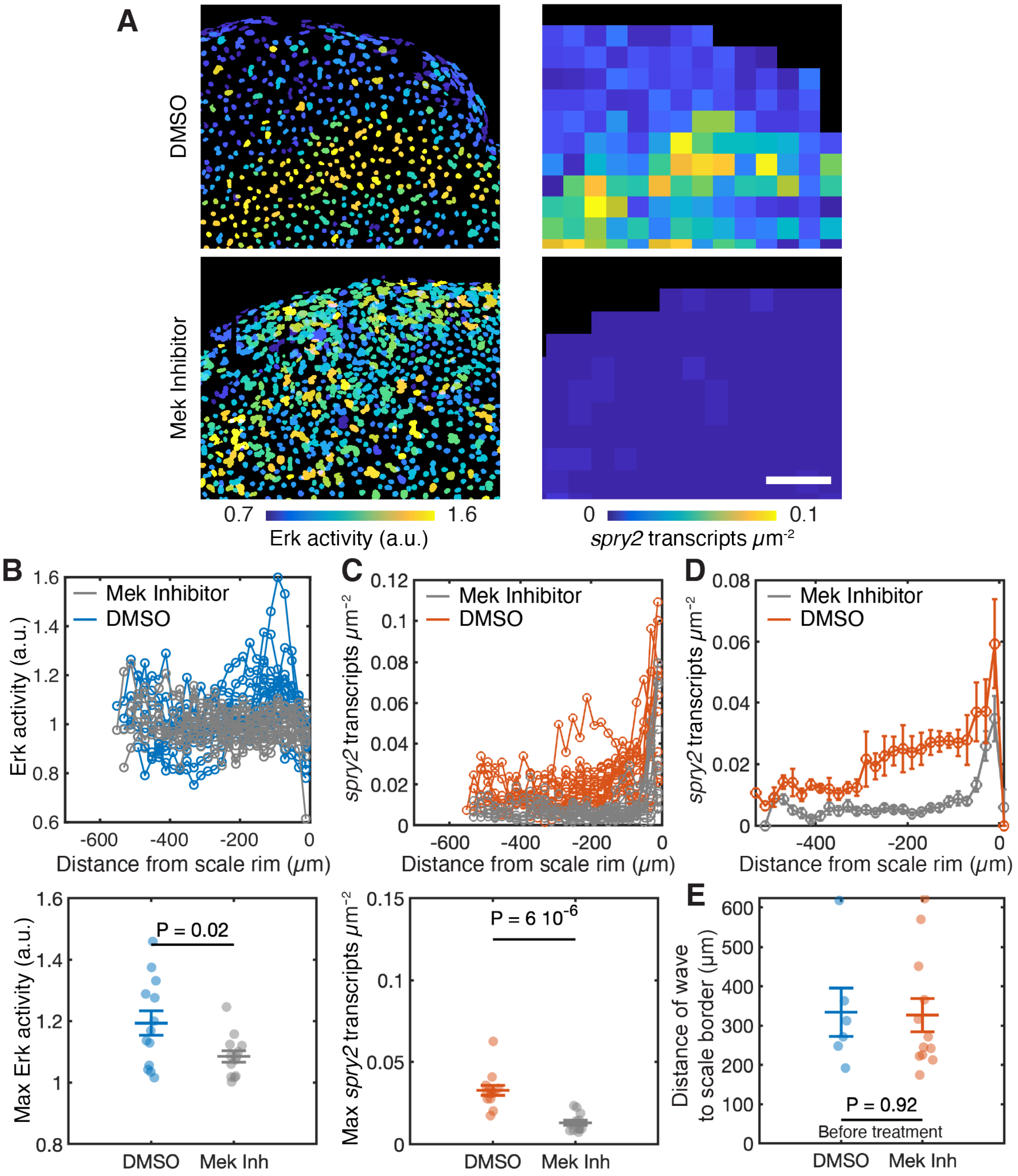


***Figure S5 –* spry2 *transcript rings are inhibited in scales treated with a Mek inhibitor.***

*A. Example of Erk activity and* spry2 *transcript density in control scales (treated with DMSO) and scales treated with the Mek inhibitor PD0325901 (10 µM). B, C. Erk activity (B, top) and* spry2 *transcript density (C, top) in control scales (treated with DMSO) and scales treated with the Mek inhibitor PD0325901 (10 µM). Amplitude of peaks of Erk activity (B, bottom) and* spry2 *transcript density (C, bottom) (with unpaired Student's t-test P-value). DMSO n = 13 scales from 4 fish; n = 2 experiments; treated with Mek inhibitor n = 13 scales from 3 fish; n = 2 experiments. D. Average* spry2 *transcript density in control scales (treated with DMSO) and scales treated with the Mek Inhibitor PD0325901 (10 µM) (same dataset as in B, C). E. Distance of the wavefront to scale border before treatment with DMSO or with the Mek inhibitor PD0325901 (10 µM) (with unpaired Student's t-test P-value; same dataset as in B). a.u: arbitrary unit. Scale bar, 100 µm.*

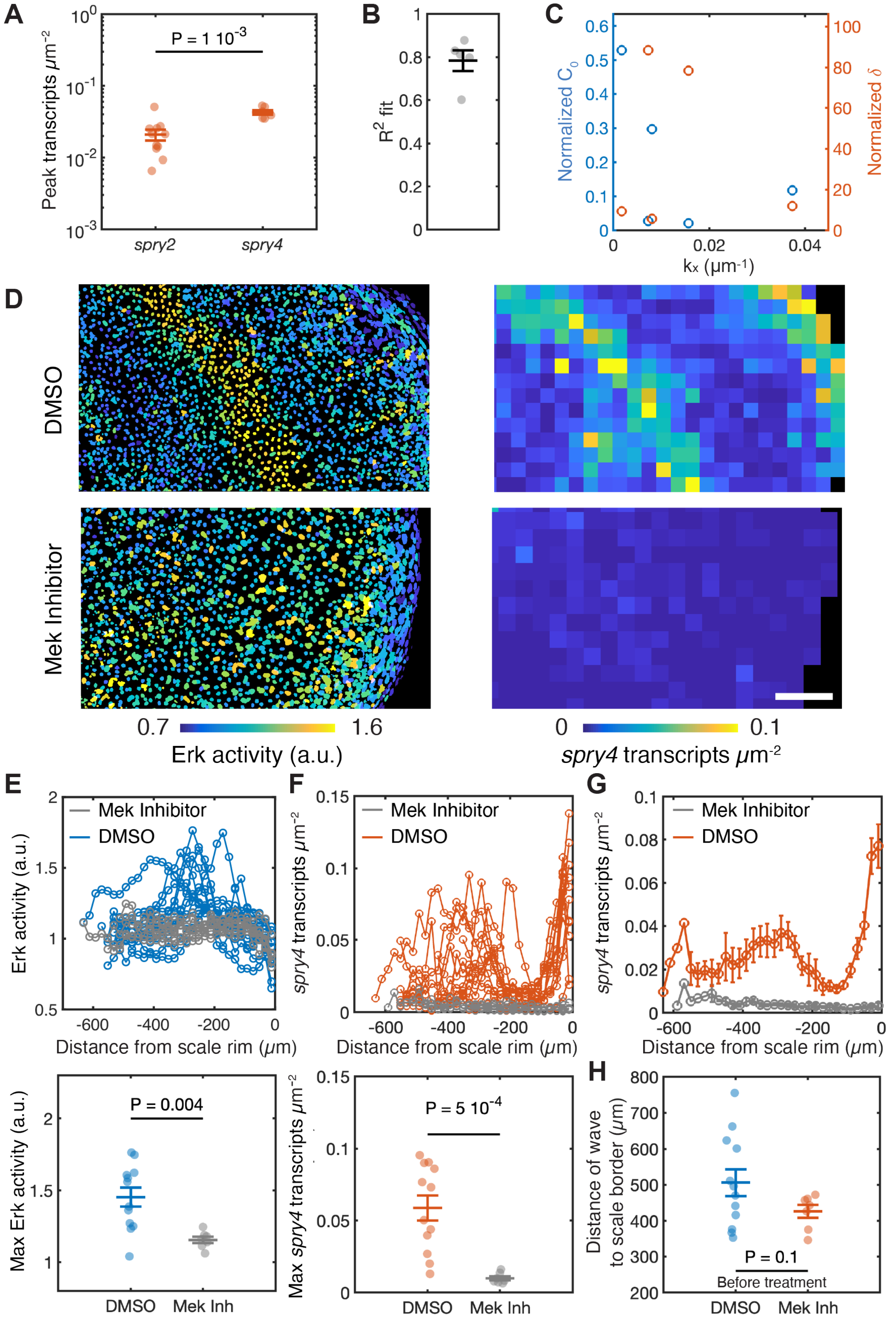


***Figure S6 - Quantification of* spry2 *and* spry4 *transcript density peaks. Fits of* spry4 *transcript profiles.* spry4 *transcript rings are inhibited in scales treated with a Mek inhibitor.***

*A. Peak amplitude of* spry2 *and* spry4 *transcript density (individual dots are individual scales from the same dataset as in Fig. 3;* spry2*: (n = 11 scales from 5 fish from 4 experiments, same dataset as Fig. 3B-D);* spry4 *(n = 6 scales from 5 fish from n = 4 experiments. same dataset as Fig. F, G); with unpaired Student's t-test P-value). The average* *peak value of* spry2 *transcript density (2 $10^{-2}$ transcripts/µ$m^2$) corresponds to ~3 transcripts per cell. The average peak value of spry4 transcript density (4 $10^{-3}$ transcripts/µ$m^2$) corresponds to ~6 transcripts per cell. B. Coefficient of determination $R^2$ values (goodness of fit) and C. fitted parameters of a fit of Erk activity-*spry4 *profiles with an ODE model in which Erk modulates* spry4 *transcript production (Methods; fitted model: $1/k_x\, dC_N/dx + C_N = C_0(1+ \delta * H(E(x)-E0))(E(x)-E0))$ in which H is the Heaviside function; dots are individual scales from the same dataset as Fig. 3F-H). D. Example of Erk activity and* spry4 *transcript density in control scales (treated with DMSO) and scales treated with the Mek inhibitor PD0325901 (10 µM). E, F. Erk activity (E, top) and* spry4 *transcript density (F, top) in control scales (treated with DMSO) and scales treated with the Mek inhibitor PD0325901 (10 µM). Amplitude of peaks of Erk activity (E, bottom) and* spry4 *transcript density (F, bottom) (with unpaired Student's t-test P-value). DMSO: n = 12 scales from 2 fish from 1 experiment; Mek inhibitor: n = 7 scales from 2 fish from 1 experiment. G. Average* spry4 *transcript density in control scales (treated with DMSO) and scales treated with the Mek inhibitor PD0325901 (10 µM) (same dataset as in E, F). H. Distance of wavefront to scale border before treatment with DMSO or with the Mek inhibitor PD0325901 (10 µM) (with unpaired Student's t-test; same dataset as in E, F) a.u: arbitrary unit. Scale bar, 100 µm.*

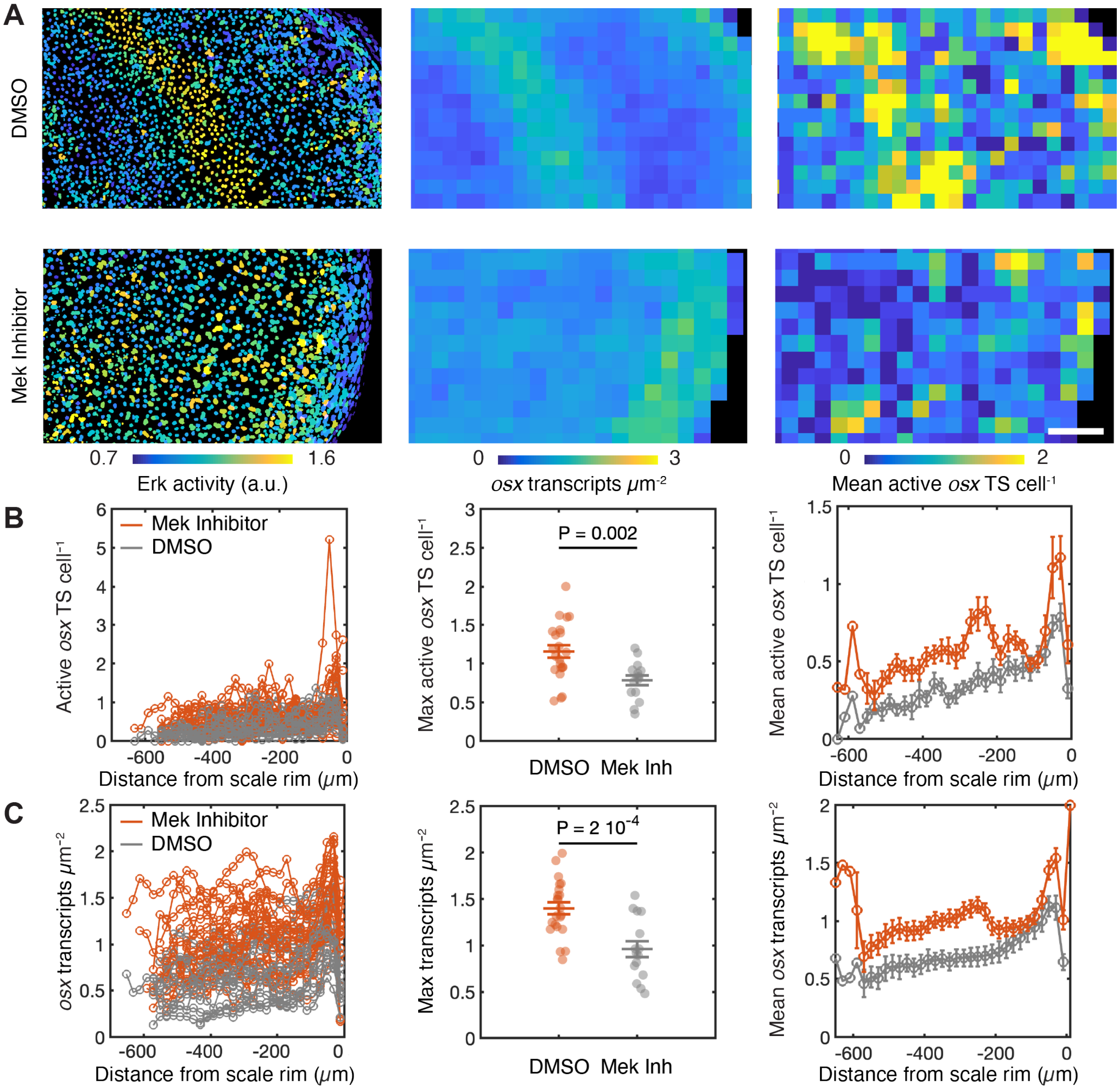


***Figure S7 –* osx *transcript rings are inhibited in scales treated with a Mek inhibitor.***

*A..Example of Erk activity (left),* osx *transcript density (middle) and number of active* osx *transcription site (TS) per cell (right; thereafter:* osx *TS per cell) in control scales (treated with DMSO) and scales treated with the Mek inhibitor PD0325901 (10 µM). B.* osx *TS per cell in control scales (treated with DMSO) and scales treated with the Mek inhibitor PD0325901 (10 µM) (left: individual; middle: peak amplitude with unpaired Student's t-test P-value; right: averages). DMSO: n = 19 scales from 4 fish from 2 experiments; Mek inhibitor n = 18 scales from 4 fish from 2 experiments. C.* osx *transcript density in control scales (treated with DMSO) and scales treated with the Mek inhibitor PD0325901 (10 µM) (left: individual; middle: peak amplitude with unpaired Student's t-test P-value; right: averages; same dataset as in B). a.u: arbitrary unit. Scale bar, 100 µm.*
